# A Lightweight Dual-Attention Neural Network for In-Situ Hyperspectral Classification of Microalgae in Underwater Monitoring Systems

**DOI:** 10.64898/2026.02.16.706104

**Authors:** Laixiang Xu, Yanyan Dong, Madineh Bijani, Yang Zhang, Xiaojie Du, Junmin Zhao

**Affiliations:** School of Information and Communication Engineering, School of Computer and Artificial Intelligence, Henan University of Urban Construction, Hainan University, Haikou 570228, China; School of Computer and Artificial Intelligence and School of Life Sciences and Engineering, International Joint Laboratory of Green and Low Carbon Water Treatment Technology and Water Resources Utilization of Henan Province, Henan University of Urban Construction, Pingdingshan 467036, China; Environmental Sciences Research Institute, Shahid Beheshti University, Tehran, Iran; State Key Laboratory of Biopharmaceutical Preparation, Chinese Academy of Sciences, Beijing, China; School of Computer and Artificial Intelligence, International Joint Laboratory of Green and Low Carbon Water Treatment Technology and Water Resources Utilization of Henan Province, Henan University of Urban Construction, Pingdingshan 467036, China; School of Computer and Artificial Intelligence, School of Life Sciences and Engineering, and School of Municipal and Environmental Engineering, Key Laboratory of Water Pollution Prevention, Control and Remediation of Henan Province, Henan University of Urban Construction, Pingdingshan 467036, China

**Keywords:** Marine ecological health, Ocean engineering application, Microalgae monitoring, Hyperspectral microscopy imaging, Fluorescence spectra, Deep learning

## Abstract

Accurate monitoring of microalgae is essential for assessing marine ecological health and preventing harmful algal blooms in ocean engineering. Current in situ identification methods often suffer from limited discriminative feature extraction and inadequate adaptation to complex underwater imaging conditions. This study introduces a lightweight dual-attention neural network, termed ANMM, designed for real-time, in situ hyperspectral classification of microalgae within integrated underwater monitoring systems. The model strengthens a deepened AlexNet backbone with multi-head latent attention (MLA) and multi-head self-attention (MSA) mechanisms, which jointly enhance local feature refinement and global spectral dependency modeling. An early-stopping strategy is further incorporated to prevent overfitting and ensure robust generalization. Evaluated on a custom dataset of field-collected fluorescence spectra, the model achieves a classification accuracy of 98.91%, outperforming several state-of-the-art deep-learning counterparts. With a compact parameter size of 16.34 M and low-latency inference on edge hardware, the system demonstrates strong potential for deployment on embedded underwater sensing platforms. This work provides a practical and efficient AI-driven solution for continuous marine microalgae monitoring, supporting advances in ocean observation technology and ecological engineering.

## 1. Introduction

Microalgae serve as vital primary producers in coastal and marine ecosystems, playing an essential role in seawater bodies and the global oceanic biogeochemical cycle (Zhao et al., 2025). Their cultivation in nearshore and offshore environments holds promise for marine bioproducts. Under illumination, marine microalgae can assimilate nutrients from aquaculture effluents, a major contributor to coastal eutrophication, and convert them into biomass, thereby aiding in nutrient pollution mitigation. However, eutrophication can also trigger the excessive proliferation of specific algal taxa, some of which produce toxins detrimental to human health, marine biodiversity, and coastal infrastructure integrity. Consequently, the accurate in-situ identification of marine microalgae is paramount for fostering beneficial species, controlling harmful algal blooms, and supporting the sustainability of ocean engineering initiatives.

Traditional manual microalgae classification in marine monitoring relies on observable traits like morphology, a process that is time-consuming and expertise-dependent. This underscores the need for efficient, automated methods suitable for coastal water monitoring. Artificial intelligence and computer vision have enabled automated image analysis and real-time species recognition. Various machine learning (Saha et al., 2025) models, including SVM and K-NN, have been applied. For instance, Xu et al. (2022) achieved 96.36% accuracy for toxic marine microalgae using K-NN, though distinguishing similar species remains difficult. Zhuo et al. (2022) used nonlinear SVM on a polarized light scattering dataset, reaching 80% accuracy with high computational cost. Anuntakarun et al. (2022) classified four algae types with 97% accuracy using random forest, requiring complex feature extraction. Similarly, Chong et al. (2024) reported 97.63% accuracy with K-NN and SVM on cultured species. Conventional ML methods typically extract handcrafted features like geometric parameters. While enhancing objectivity, performance depends heavily on feature engineering and lacks generalizability across varying marine growth stages and imaging conditions, limiting accuracy in real-world offshore applications.

To overcome these limitations, deep learning (Slaughter et al., 2025), particularly convolutional neural networks (CNNs), has become a mainstream solution for algae image classification. CNNs fundamentally automate feature extraction, eliminating the reliance on manual feature engineering and conserving significant human and computational resources. By learning complex, hierarchical feature representations directly from images, CNNs typically achieve superior accuracy in recognizing algae within complex backgrounds and diverse morphologies. Moreover, well-trained CNN models exhibit strong generalization, maintaining high performance on unseen samples. For instance, recent studies report accuracies from 87% to 91.20% (Liu et al., 2024) and 95.67% (Shen et al., 2025) for microalgae classification. Higher performance has been demonstrated in specific contexts: Yang et al. (2023) achieved 98.0% recall for plankton identification, Yuan et al. (2023) reached 99.87% accuracy in algal bloom monitoring, and Gaur et al. (2022) obtained 99.1% accuracy for blue-green algae recognition.

Despite these advances, deep learning approaches for microalgae recognition still face key challenges: inadequate extraction of multi-scale spectral features, a complexity-efficiency trade-off that limits real-time application, and poor generalization in data-scarce monitoring scenarios. To overcome these issues, this paper introduces a lightweight enhanced AlexNet integrated with multi-head attention mechanisms for hyperspectral microalgae classification. The model strengthens both local and global feature representation while maintaining deployability on common hardware, bridging robust feature learning with operational efficiency for practical in-situ analysis. The main contributions of this work are as follows:

1. Incorporating multi-head latent attention and multi-head self-attention modules to enhance the joint modeling of local details and global features.
2. Integrating an early stopping mechanism with dynamic learning rate scheduling to improve training efficiency and model generalization.
3. On a custom-built dataset, the proposed model achieved 98.91% classification accuracy, outperforming several mainstream deep learning models and offering a lightweight, high-accuracy solution for real-time microalgae monitoring and classification.

The remaining part of this article is as follows: Part 2 offers the microalgae spectral data collection setup. Part 3 mainly introduces the proposed method. Part 4 reports experimental results and analysis. Part 5 provides conclusions and future research directions.

## 2. Microalgae spectral data collection

We used a self-built multi-mode hyperspectral microscope imaging equipment to gather spectral data of four microalgae species in Dongpo Lake, Hainan University. This system integrates two illumination modes: layer light and wide-field light, mainly consisting of multi-mode illumination units and multi-channel optical imaging units. The lighting unit is responsible for the generation and regulation of layer light and wide-field light, covering the design of corresponding optical path structures and driving circuits. The imaging unit includes a hyperspectral camera and a dual-channel microscope. Synchronous capture of fluorescence, scattering, and hyperspectral signals is accomplished by fine-tuning the filter arrangement. The system’s operation is depicted in Fig. 1. To enhance the integration and applicability of the system, we have designed an optical imaging module (Fig.1) that can simultaneously capture fluorescence microscopy images, scattering microscopy images, and hyperspectral data. This module uses a flat field infinite distance objective lens (Fig. 1(15)) to image the sample, achieving a collimated state of the sample beam. Subsequently, it utilizes three detection channels to obtain optical information in parallel. To achieve synchronous acquisition, a 90/10 beam splitter (Fig. 1(14)) is integrated into the cage cube at the back of the objective lens, which reflects 90% of the light to the spatial imaging channel and transmits 10% to the spectral detection channel. According to the principle of microscope optics, we have equipped a tube lens behind the infinity objective lens (Fig. 1(13)) to focus the optical signal. Subsequently, the beam was split into two paths using a 50/50 beam splitter (Fig.1(11)): one path was reflected and directly entered the CMOS camera for scattering microscopy imaging. Another is to install a 450 nm filter in front of the CMOS camera (Fig.1(12)) to filter out excitation light and achieve fluorescence microscopy imaging.

**Fig. 1:**
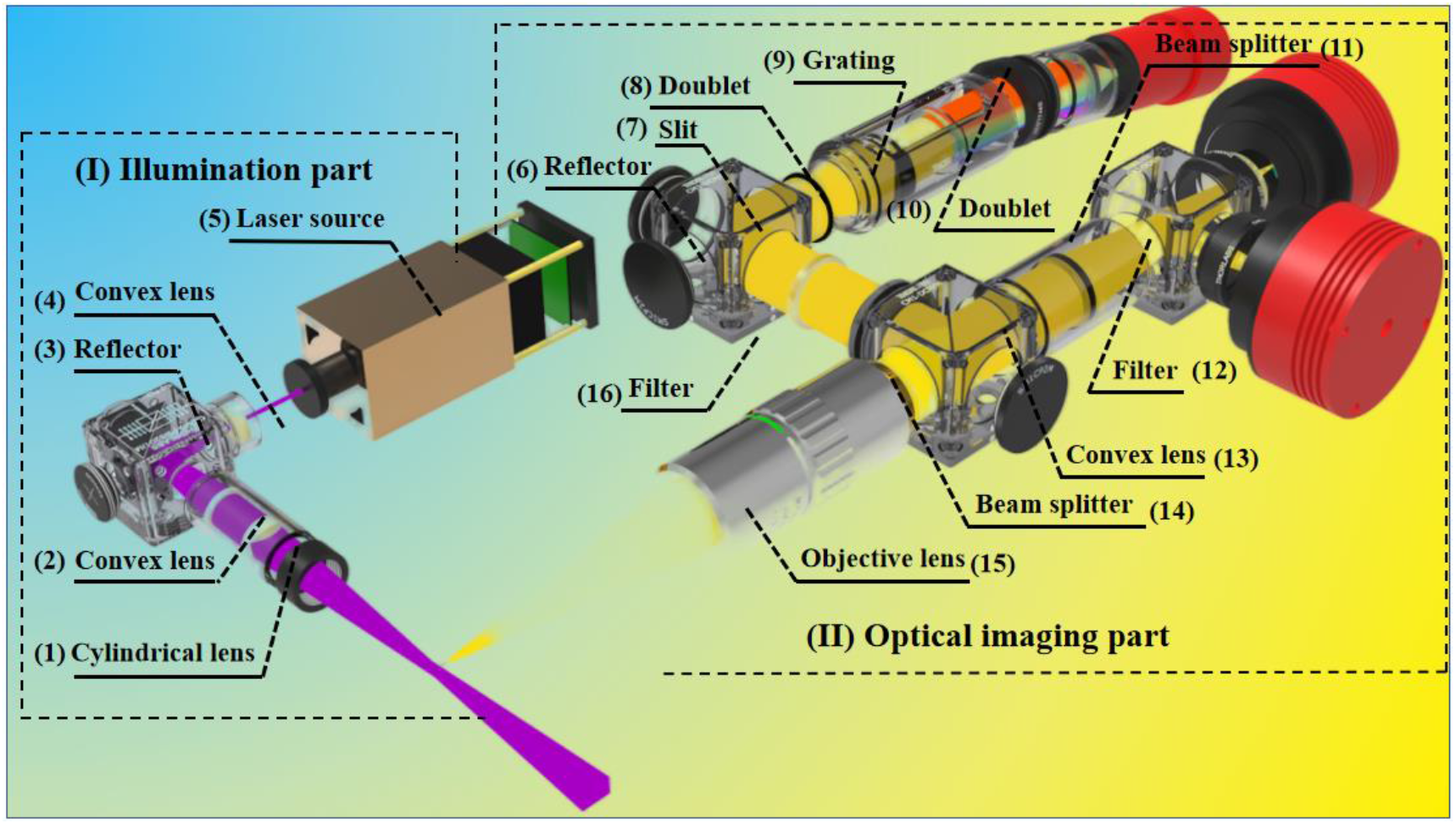
Lighting and optical imaging architecture.

The core of a spectrometer is the spectroscopic system, which functions to spatially separate incident light by wavelength. The system consists of three main components: a collimation unit, a dispersion unit, and a photoelectric sensing unit. The collimation unit consists of an imaging lens and a slit, with the incident slit set at the focal plane position of the collimation objective lens. After passing through the imaging lens, the light beam converges at the slit and only retains the light along the direction of the slit. To suppress excitation light, a 450 nm filter is installed in front of the slit, allowing only fluorescence signals to pass through. Subsequently, the fluorescent signal is guided to the grating through a pair of laminated lenses, where the composite light is decomposed into multiple monochromatic light beams. Finally, another pair of laminated lenses is used for color difference correction, and the separated monochromatic light is focused onto the rear photoelectric sensor and converted into an electrical signal output.

To cope with the complex and ever-changing working conditions underwater, we have developed a sealed cabin structure that enables the entire system to be installed in a distributed layout in the underwater environment (as shown in Fig. 2). The core optical part consists of a hyperspectral camera, multi-channel microscope, and sheet-like light source, which are firmly installed in the waterproof compartment of the cabin with the help of metal brackets and obtain optical signals through specially reserved tempered glass windows. After completing airtightness, strength, and rigidity tests, the system has been proven to withstand pressure up to 50 meters deep and ensure continuous and stable operation of internal optical equipment.

**Fig. 2:**
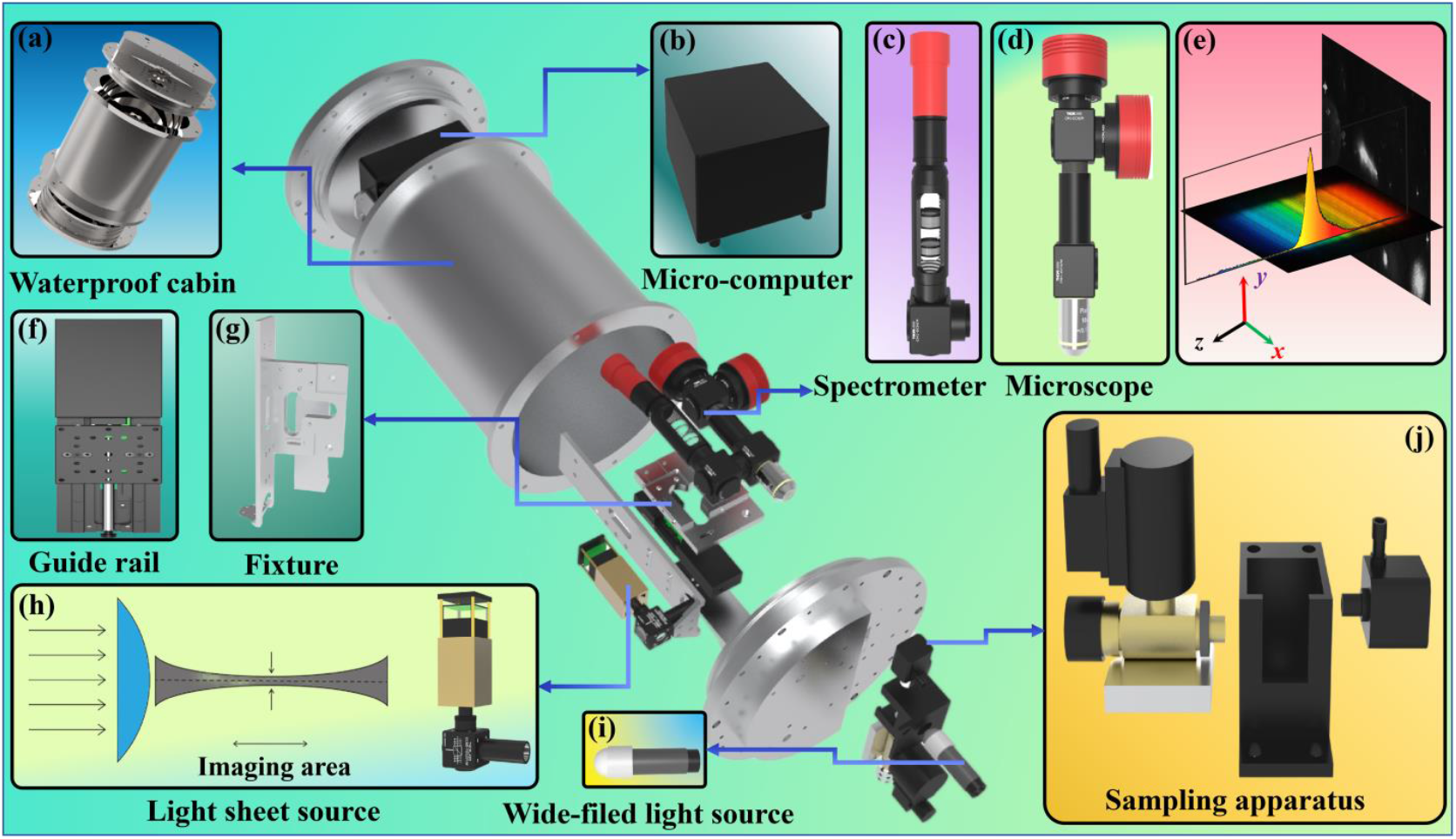
(a) Watertight pressure housing; (b) Microcomputer unit; (c) Hyperspectral camera; (d) Microscopic imaging module; (e) 3D schematic view of the imaging results; (f) High-precision linear guide rail; (g) Mounting fixture for an optical system; (h) Light sheet illumination source with an optical path diagram for a cylindrical lens; (i) Broad-field illumination source; (j) Sealed sample chamber with an electronic water valve and pump.

The underwater sealed cabin adopts a sandwich layout, including a main cabin and two end caps. There are two waterproof cables installed on the front cover, which are connected through a waterproof joint. One of them is responsible for delivering power from the shore-based power source to the underwater system, ensuring the stable operation of electronic components. The system is equipped with a multi-channel voltage conversion module, which can rectify and filter AC power while outputting different DC voltages such as 12V and 5V to meet the rated power requirements of various electronic devices. The other cable follows the Ethernet transmission line standard, which not only enables communication between the water control terminal and the underwater cabin but also ensures stable and reliable data exchange.

The major peak of the microalgae fluorescence spectrum is mostly dispersed between 500 and 800 nm, with peaks centered between 650 and 700 nm. Based on this, we truncated the spectrum and only retained the light intensity information within the 500 nm–800 nm band, extracting a total of 811 pixels. Each type of microalgae contains 100 spectral curves, with a total of 400 curves for the four types, each curve containing 811 pixel values. Due to the fact that the main peak of microalgae fluorescence signals is located in this band, and there are significant differences in spectral characteristics among different species in this range, the fluorescence spectrum image is segmented by wavelength, and only this band is retained to reduce the interference of noise in other regions in the subsequent recognition process. Subsequently, we set a threshold to perform binary processing on the image: the parts below the threshold are treated as background and set to 0, while the parts above the threshold are treated as effective signals and set to 1. Then, multiple discrete effective spectral intervals are extracted using the connected domain detection method, and the original spectra within each interval are superimposed along the vertical direction of the slit to finally calculate the single-molecule fluorescence spectra of microalgae cells in each interval.

Due to the high dimensionality of one-dimensional spectral data, direct use in training can lead to the loss of important information and suboptimal recognition performance. Since deep learning algorithms excel at processing two-dimensional images, we propose converting the one-dimensional fluorescence spectra into 2D images to improve analytical efficiency. Specifically, a central sorting method is employed for this transformation. This method positions the median value of the spectral sequence at the center of the image and arranges the remaining data points in a spiral pattern outward. For computational consistency, each resulting image is stored as a 32×32 matrix, with zero padding applied where necessary. This approach effectively enhances data dimensionality while preserving the intrinsic wavelength-order relationships of the original spectra.

A hyperspectral microscopy imaging system was used to collect one-dimensional fluorescence spectral sequences from five microalgae species at Dongpo Lake, Hainan University, and Daya Bay Nuclear Power Station, Shenzhen. Each sequence was converted into a two-dimensional image via the central sorting method described above, preserving spectral structure and providing a richer input format for convolutional neural networks. The complete preprocessing workflow is illustrated in Fig. 3.

**Fig. 3:**
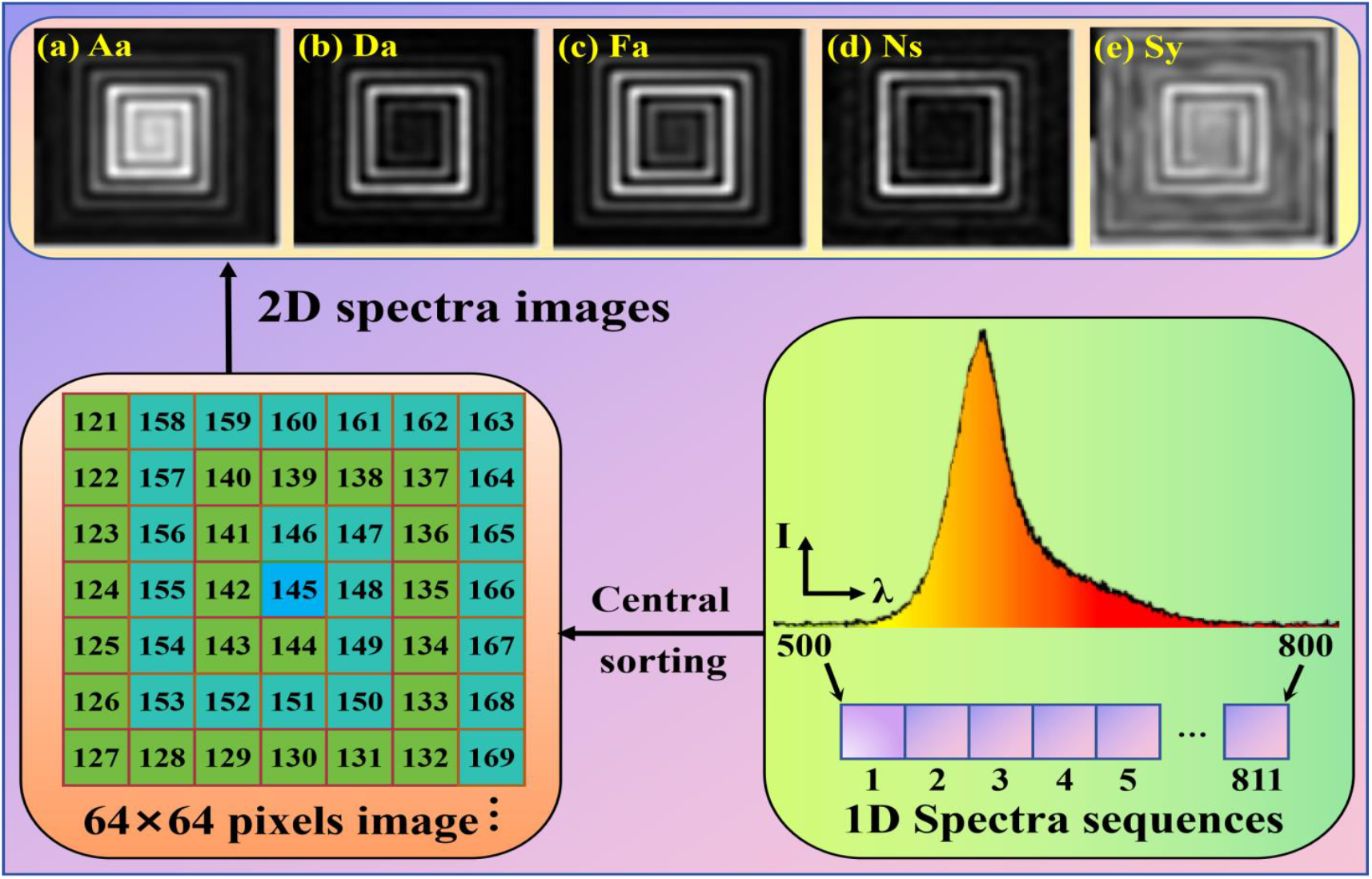
The process of converting one-dimensional spectral data into two-dimensional images.

## 3. Proposed methods

### 3.1 Improved AlexNet

In this study, we adopt AlexNet as the foundational architecture for classifying microalgae using two-dimensional center-sorted spectral representations. These representations display intricate, multi-scale patterns characterized by pronounced inter-band dependencies. Despite its effectiveness in general image recognition, the standard AlexNet suffers from limited network depth and insufficient feature abstraction, which constrains its capacity to capture the hierarchical spectral structures essential for precise taxonomic discrimination.

To improve feature extraction, an extra convolutional layer is inserted after the fourth convolutional layer. The first three layers capture basic local spectral gradients, while the fourth begins modeling intermediate patterns. The added fifth layer integrates cross-band correlations, transforming spectral responses into discriminative biochemical representations.

The improved model comprises an input layer, six convolutional layers, three pooling layers, three fully-connected layers, and an output layer. Its architecture is detailed in Fig. 4.

**Fig. 4:**
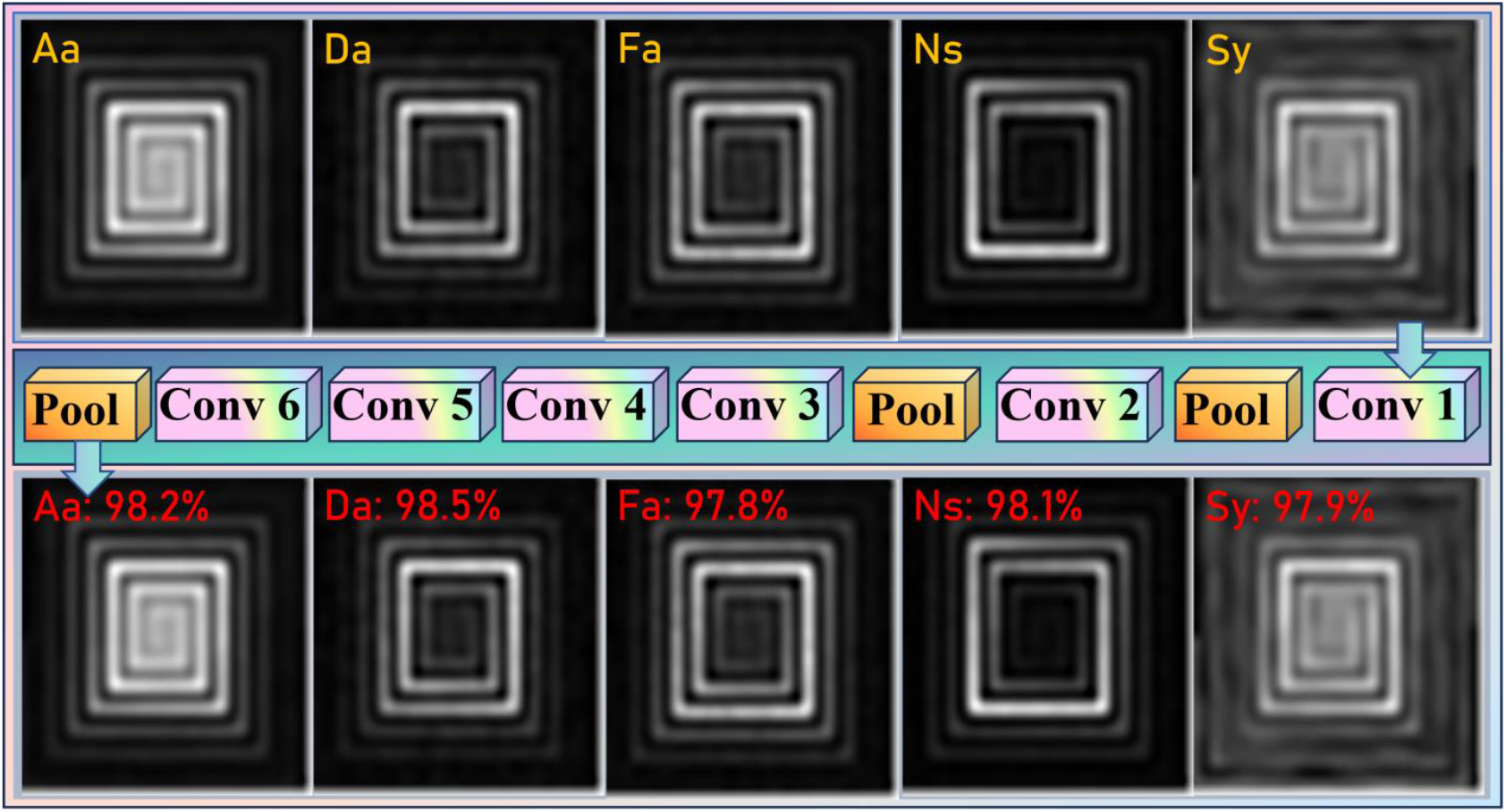
The Network Structure of Improved AlexNet

The convolution operation serves as the core of the network. It scans the input two-dimensional spectral image using a set of learnable filters (kernels). Each filter specializes in extracting a specific type of local spectral pattern (e.g., an absorption peak or a fluorescence shoulder at specific wavelengths). This design of local receptive fields, combined with weight sharing, significantly reduces the number of model parameters and provides the network with a foundational translation invariance. The convolution operation can be expressed as follows:

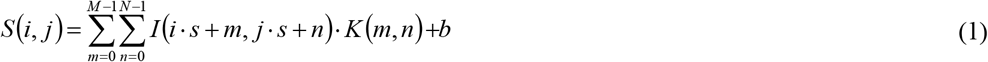

In this formulation, I is the input feature map, and K is a convolution kernel of dimensions M×N×C×D, where the first two dimensions define its spatial size, C is input channels, and D is output channels. The convolution stride is s, b is the bias, while m and n are the row and column indices within the kernel.

The pooling layer follows the convolutional layers, performing down-sampling on the feature maps. It compresses the spatial dimensions while preserving the most significant activations. This operation not only reduces computational load for subsequent layers and accelerates training but also introduces a degree of robustness to minor shifts and deformations in the spectral signal. This model uses the maximum pooling approach. Its mathematical expression can be described as:

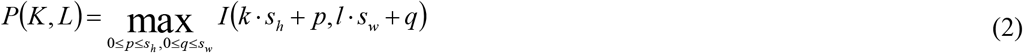

In which, *s*_*h*_ and *s*_*w*_ are the height and width of the pooled window, offsets of rows and columns in the pooled window are represented by p and q, but the row and column indexes of the pooled feature map are represented by k and l, and after pooling. *P*(*K, L*) is the value at position (*k,l*) on after pooling.

Three or so rounds of convolution and pooling operations are followed by a flattening operation on the feature map which then is fed to fully connected layers. These layers combine the locally prominent features with semantic representations that are globally coherent and therefore allowing eventual classification decisions. The affine transformation is done between the high-dimensional features to the output class space by the fully connected layers mapping the features to the output classes as a weight matrix and a bias vector. The mathematical formulation of a fully connected layer can be expressed as follows:

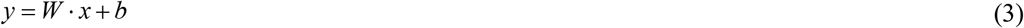

in which, y denotes the vector of output scores, which is derived from the flattened output eigenvector x through a linear projection defined by weight matrix W and offset vector b.

These scores are subsequently converted into a probabilistic form by a Softmax classifier. The mathematical formulation of the Softmax operation is provided below:

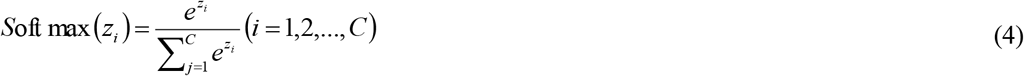

where z represents the classification score vector output by the full connection layer, dimension is C, *z*_*i*_ denotes the score value of the ith category, and the output means the prediction probability of the ith category.

in which, z is the C-dimensional classification score vector produced by the fully-connected layer. The elemen *z*_*i*_ corresponds to the score for the ith category, and the output signifies the predicted probability for that category.

### 3.2 Multi-head latent attention

The improved AlexNet improves intermediate feature representation, yet its reliance on convolutional layers with localized receptive fields still limits the modeling of long-range spectral interactions. When processing 2D center-sorted spectral images, this locality constraint leads to three primary shortcomings, such as inadequate modeling of cross-region spectral dependencies, diminished sensitivity to subtle spectral signatures, and increased vulnerability to noise interference during feature selection. Consequently, features extracted by convolutional layers alone often retain substantial redundancy and insufficiently emphasize discriminative spectral patterns.

To overcome these constraints, the MLA mechanism is introduced after the fifth convolutional layer. At this stage, the feature maps encapsulate high-level spectral abstractions while retaining the spatial resolution necessary to preserve local detail. Operating directly on these refined features, the MLA module concentrates on semantically salient spectral regions and minimizes interference from low-level redundancies. By modeling interactions across both channel and spatial dimensions, it strengthens the model’s ability to emphasize subtle yet discriminative spectral signatures. The Architecture of proposed MLA module is presented in Fig. 5.

**Fig. 5:**
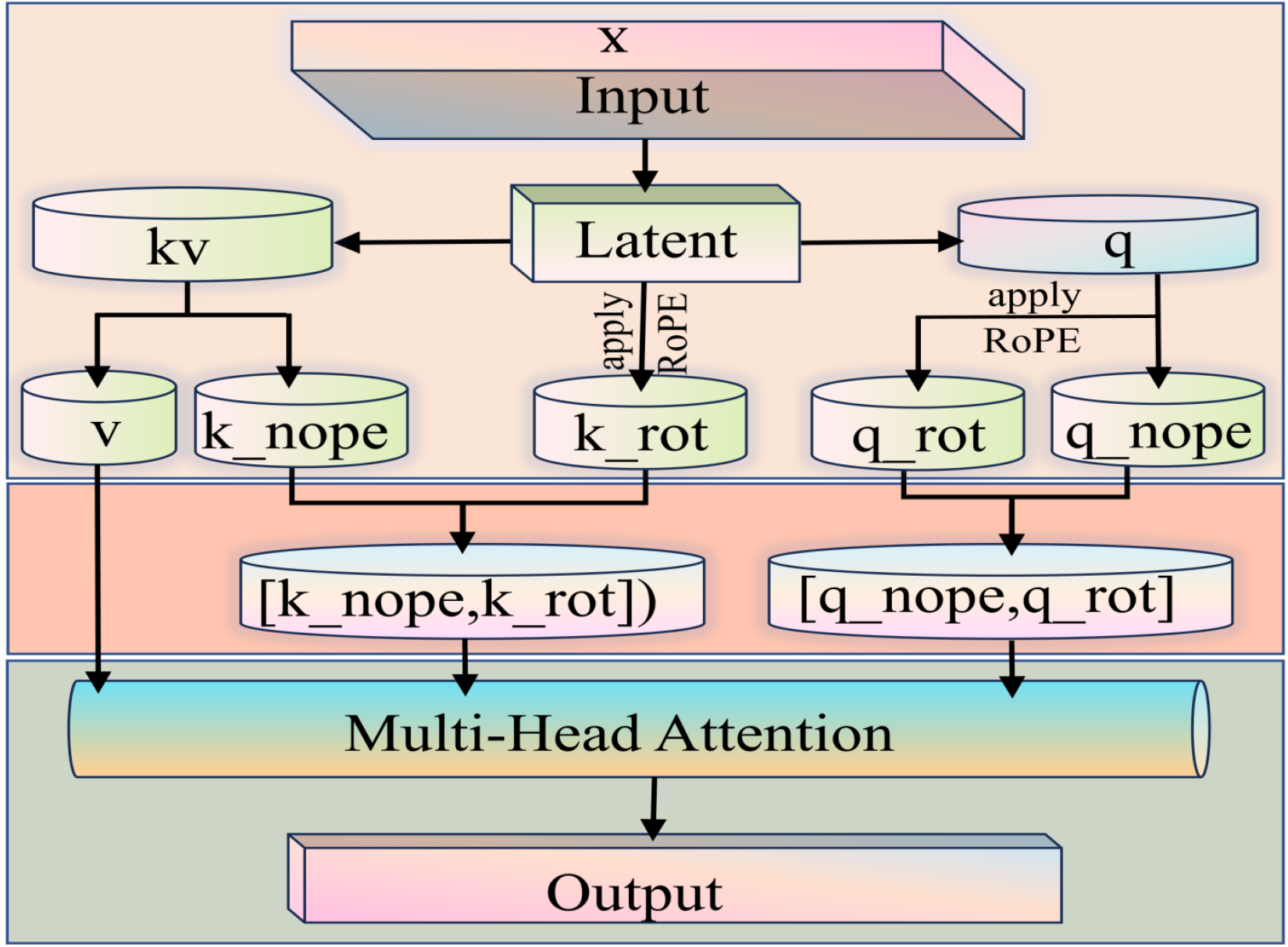
Architecture of the proposed MLA module. The inputs are defined as the k, q, and v. In the latent processing section, the q and the combined kv are received, and their forward path to the next component is illustrated by directional arrows.

The essence of MLA lies in compression of key and values with low rank in order to shrink the cache. Their expressions are computed in the following way:

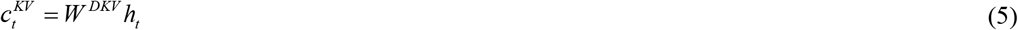

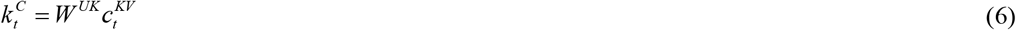

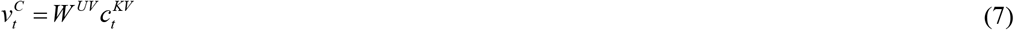

Where 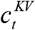 is the compressed potential vector of key and value, *W*^*DKV*^ means the down projection matrix, *W*^*UK*^ and *W*^*UV*^ are the up projection matrix of key and value respectively, and MLA only needs to cache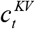.

Despite offering no reduction to the KV cache, low-rank compression is applied to the query to decrease active memory usage during training. The procedure is defined as follows:

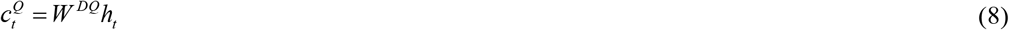

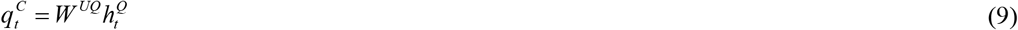

where the query’s compressed potential vector is introduced, and where *W* ^*DQ*^ and *W*^*UQ*^ are respectively designated as the lower and upper projection matrices for this vector.

### 3.3 Multi-head self-attention

While the MLA module excels at extracting fine-grained local spectral attributes, it does not sufficiently model global contextual relationships across the spectral domain. This shortcoming becomes pronounced when classifying 2D center-sorted spectral data, where inter-class spectral similarities and environmental noise can degrade discrimination performance. To address this, we introduce MSA module following the last convolutional layer, enabling the network to integrate non-local spectral dependencies.

At this stage, feature maps contain high-level semantic representations, allowing MSA to capture long-range dependencies across the spectral domain, suppress noise, and enhance globally distinctive patterns. Together, MLA and MSA form a complementary dual-attention framework: MLA preserves local spectral signatures, while MSA integrates broader contextual relationships. This synergy improves both discriminative power and model robustness, leading to higher classification accuracy. The hierarchical design aligns with the multi-scale nature of spectral data and promotes interpretable feature learning. The structure of the MSA module is shown in Fig. 6.

**Fig. 6:**
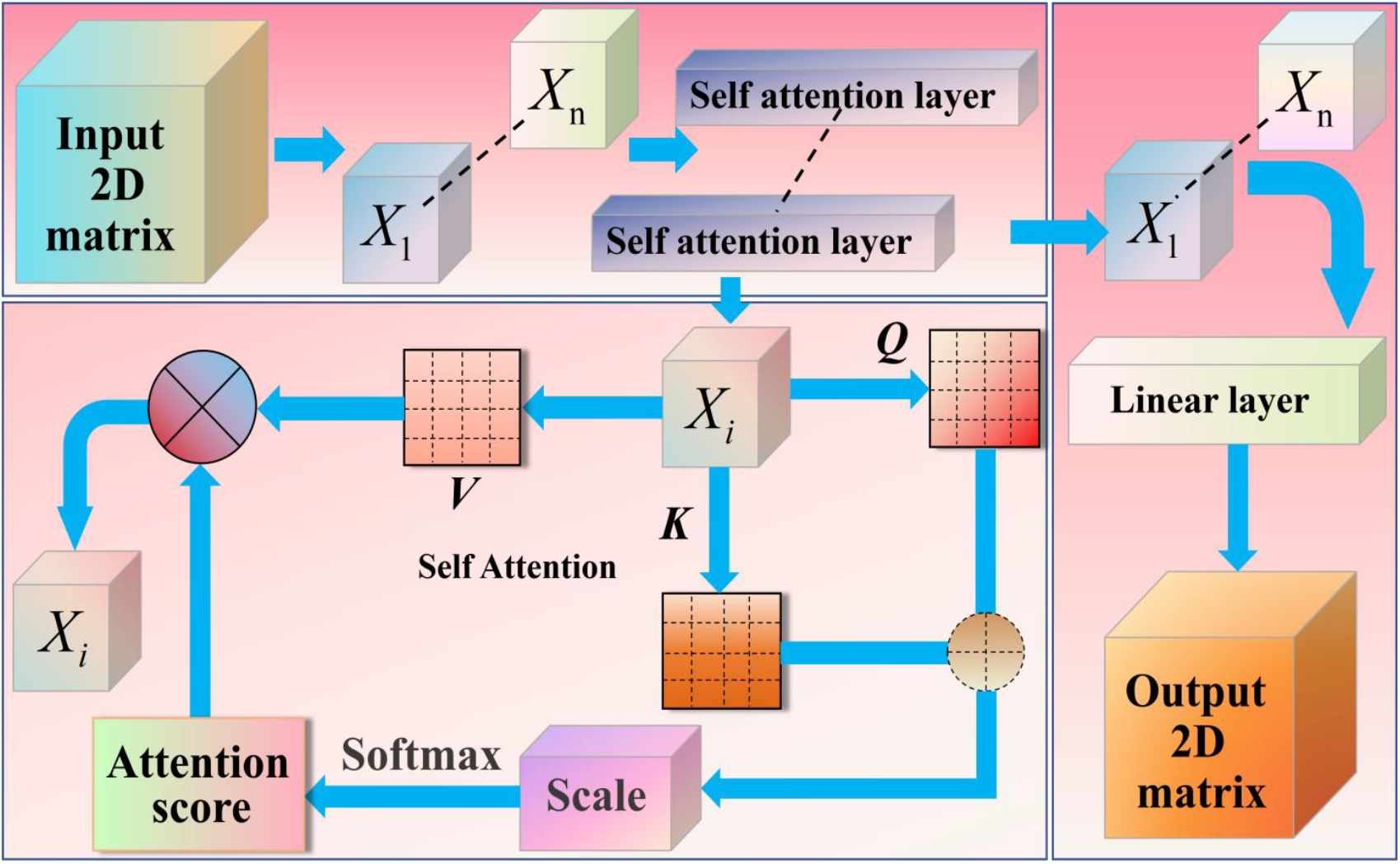
Flowchart of the proposed MSA mechanism. The model processes input signals through a self-attention module to generate the output.

One of its steps is the transformation of inputs controlling the mechanism and linear projection. Linear projection is not only useful in the mapping of the input data in the various representation spaces by the model, but also in enabling the model to learn the dynamics of focusing on various elements of the input data depending on the requirements of the task at hand. Once the input is turned into a sequence, it gets mapped to a query,key and value m. This step is obtained via multiplying matrices by weight matrices,and each of the matrices is a learned - weight e. Its calculation method is given by:

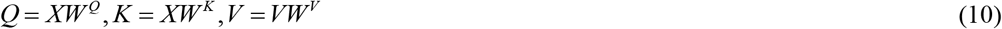

where X is the input sequence, *W* ^*Q*^ and *W* ^*K*^ and *W* ^*V*^ represent learnable parameters.

In multi-head attention, actually there will be many linear transformations of query, key and value, and their weight matrices are separately. In this way, the input vector will split into many different subspaces, and every subspace will be implemented with self-attention operation. Its mathematical expression can be described as.

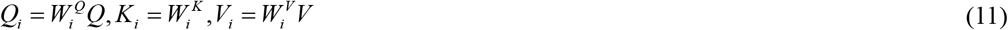

Where 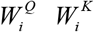 and 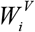 are the transformation matrices of the query, key, and value of the ith header, respectively.

At this stage, features represent high-level abstractions. The MSA module enables each position in the feature map to interact with all other positions, modeling the global contextual relationships across the entire spectral range. This is vital for distinguishing species with similar local peaks but different overall spectral shapes. The scaled dot-product attention mechanism is employed across multiple heads to learn diverse global dependencies:

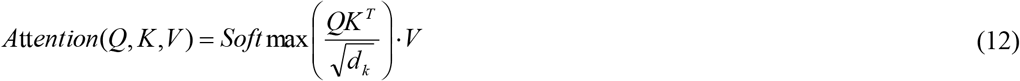

The function of the scale factor 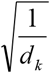 is to avoid large dot product results and make the dimensional normalization results tend to 0 or 1.

MSA employs h distinct linear projection weights for Q, K, and V to obtain h different attention heads. Their outputs are merged and projected to the final dimension. The computation is given by:

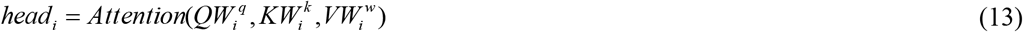

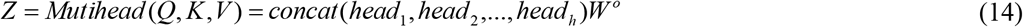

Where *head*_*i*_ s the output of the i-th attention head, with concat indicating matrix splicing and *W*^*O*^ being the output layer’s weight matrix, respectively.

To improve training efficiency and generalization, an early stopping mechanism alongside a learning rate scheduler is adopted. Training When the validation performance stops improving over a specified number of epochs, it halts training and the scheduler decreases the learning rate, effectively stabilizing the convergence and discouraging overfitting. All these strategies encourage the efficient and computationally strong model performance. The general layout of the suggested framework of microalgae classification is depicted in Fig. 7.

**Fig. 7:**
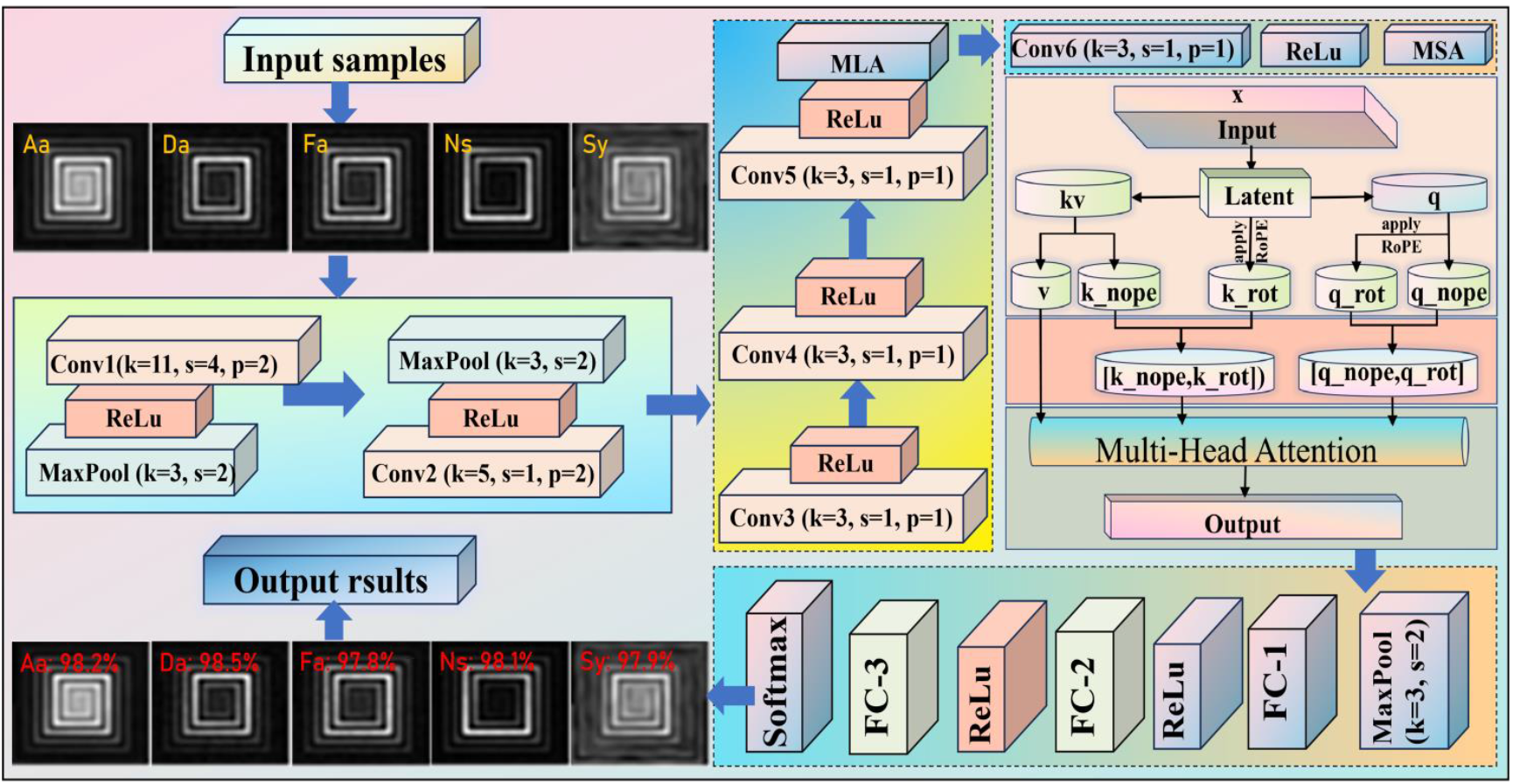
The overall recognition framework of algae classification model.

The 2D spectral image is first processed by the initial five convolutional and two pooling layers of the enhanced AlexNet. The MLA module then operates on the resulting feature maps, dynamically emphasizing discriminative local spectral bands and suppressing noise through its latent attention mechanism. Subsequently, the MSA module integrates global contextual information from the features produced by the sixth convolutional layer, modeling relationships across the entire spectral representation. The refined features are finally passed through the fully-connected layers and a Softmax classifier to output the predicted microalgae species probabilities.

## 4. Experimental results and analysis

### 4.1 Experimental setup

All experiments were performed using a workstation with the use of an AMD Ryzen 7 4800H processor, 32 GB RAM, and NVIDIA GeForce GTX 1650 Ti (4GB of VRAM) graphics card. Applications (software stack) Windows 10 version (64-bit), Python 3.12, PyTorch 2.3 deep learning architecture, CUDA 11.8, cuDNN 8.7 to accelerate the software with the help of a graphic card.

### 4.2 Data description

Given the limited number of samples initially collected, data augmentation techniques, including rotation, scaling, flipping, and cropping, were applied to expand the dataset. This process enriches the feature variability available for training, which is crucial for enabling the deep learning model to learn robust and generalizable representations from a larger effective sample size. By simulating natural variations in imaging conditions and microalgae presentation, these transformations help mitigate overfitting and improve the model’s ability to handle real-world diversity. The distribution of original and augmented samples for each microalgae category is summarized in Table 1.

**Table 1:** Description of Algae Spectral Data.

| Category | Abbreviation naming | Original quantity | Enhance quantity | Total number |
| --- | --- | --- | --- | --- |
| Aeruginosa | Aa | 100 | 2200 | 2300 |
| Dunaliella salina | Da | 100 | 2200 | 2300 |
| Flat algae | Fa | 100 | 2200 | 2300 |
| Nannochloropsis | Ns | 100 | 2200 | 2300 |
| Synechocystis | Sy | 100 | 2200 | 2300 |

Using a hyperspectral microscopy imaging system, one-dimensional fluorescence spectral sequences were collected from five microalgae species at Dongpo Lake, Hainan University, and Daya Bay Nuclear Power Station, Shenzhen. Following data augmentation, a total of 11,500 two-dimensional fluorescence spectral images were generated for subsequent analysis.

To balance experimental efficiency with reliable evaluation, the dataset was partitioned into training, validation, and test subsets following a 3:1:1 ratio. The final dataset comprises 6,900 training, 2,300 validation, and 2,300 test images. The detailed sample distribution across the five microalgae categories is provided in Table 2.

**Table 2:**
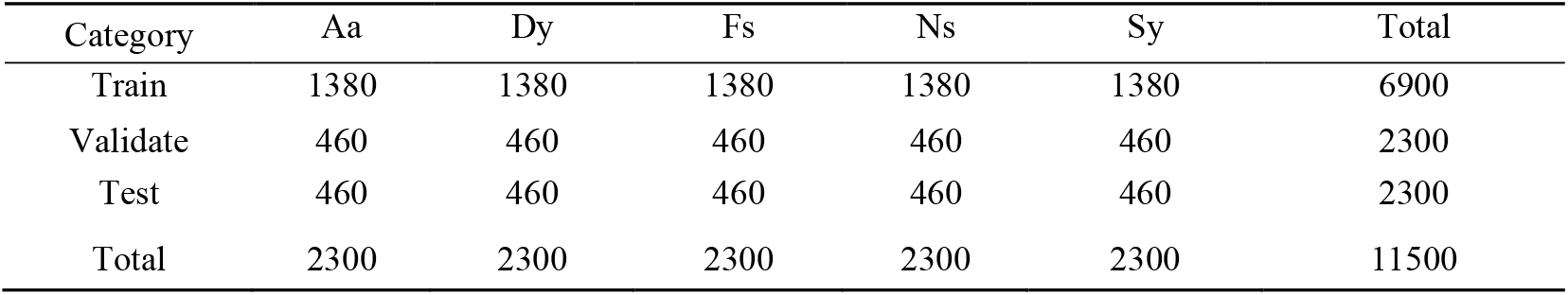
Data distribution of each type of algae.

| Category | Aa | Dy | Fs | Ns | Sy | Total |
| --- | --- | --- | --- | --- | --- | --- |
| Train | 1380 | 1380 | 1380 | 1380 | 1380 | 6900 |
| Validate | 460 | 460 | 460 | 460 | 460 | 2300 |
| Test | 460 | 460 | 460 | 460 | 460 | 2300 |
| Total | 2300 | 2300 | 2300 | 2300 | 2300 | 11500 |

### 4.3 Train and test results

To compare the training performance of different models, we present the changes in training and validation accuracy loss values in Fig. 8.

**Fig. 8:**
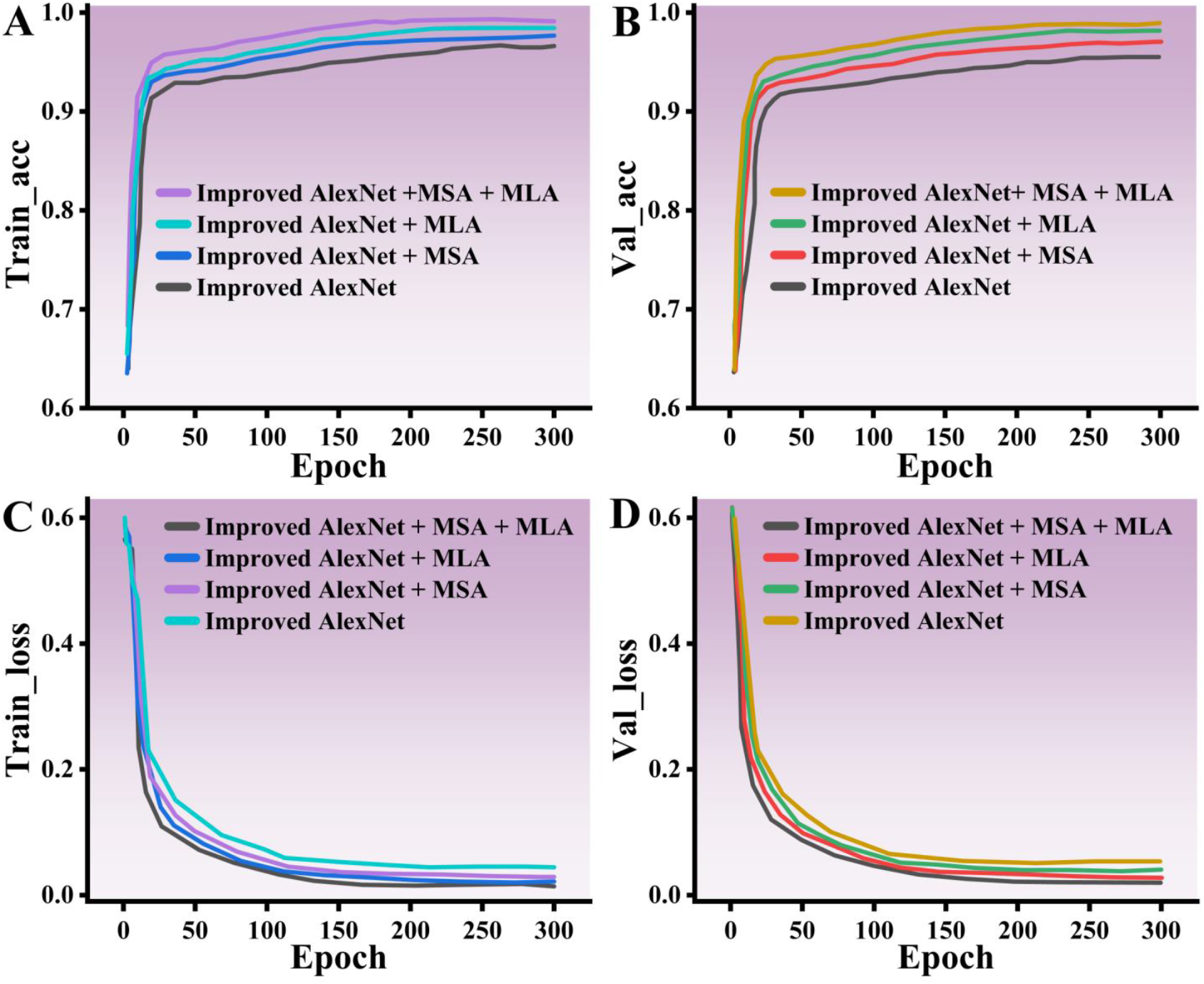
The accuracy and loss variation curves for the four methods. Subfigures A and B present the training and validation accuracy, while C and D depict the corresponding training and validation loss.

As shown in Fig. 8, the training and validation accuracy of the model progressively stabilizes with increasing epochs, showing minimal fluctuation after 200 iterations. Among the compared architectures, Improved AlexNet with MLA achieves higher accuracy than its counterpart with MSA, indicating that the MLA mechanism alone contributes more substantially to performance improvement. The proposed model, which integrates both MLA and MSA, attains the lowest loss and highest accuracy among all four configurations. This superior result stems from the synergistic interaction between the two attention mechanisms: MLA enhances the capture of locally discriminative spectral details, while MSA strengthens the modeling of global contextual dependencies across spectral bands. The stable convergence of the accuracy and loss curves confirms the model’s strong feature extraction capability and training stability. By jointly leveraging local refinement and global integration, the combined attention design enables more robust and accurate recognition.

Recall, Precision, Specificity, F1-score, and Accuracy are standard metrics for evaluating classification models (Zarboubi et al., 2025). Their definitions are as follows: Recall is the proportion of actual positives correctly identified. Precision is the fraction of predicted positives that are true positives. Specificity measures the proportion of actual negatives correctly identified. The F1-score is the harmonic mean of precision and recall, balancing both. Accuracy is the overall percentage of correctly classified samples. These metrics are formally expressed below:

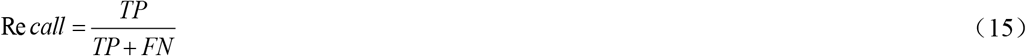

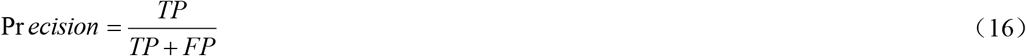

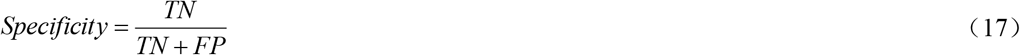

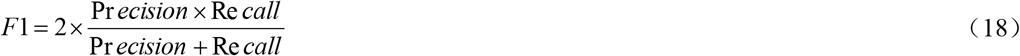

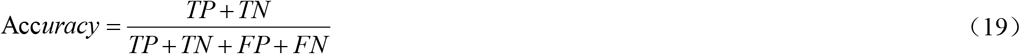

Radar charts offer an efficient way to visualize multidimensional performance metrics within a unified coordinate framework. They allow for clear comparative analysis of the relative strengths and weaknesses of different models across various microalgae classification tasks. In this work, radar charts are used to display model performance across five key evaluation metrics. As shown in Fig. 9, this visualization supports a systematic and comprehensive comparison of the diagnostic ability of each model, highlighting how well they balance different aspects of classification performance.

**Fig. 9:**
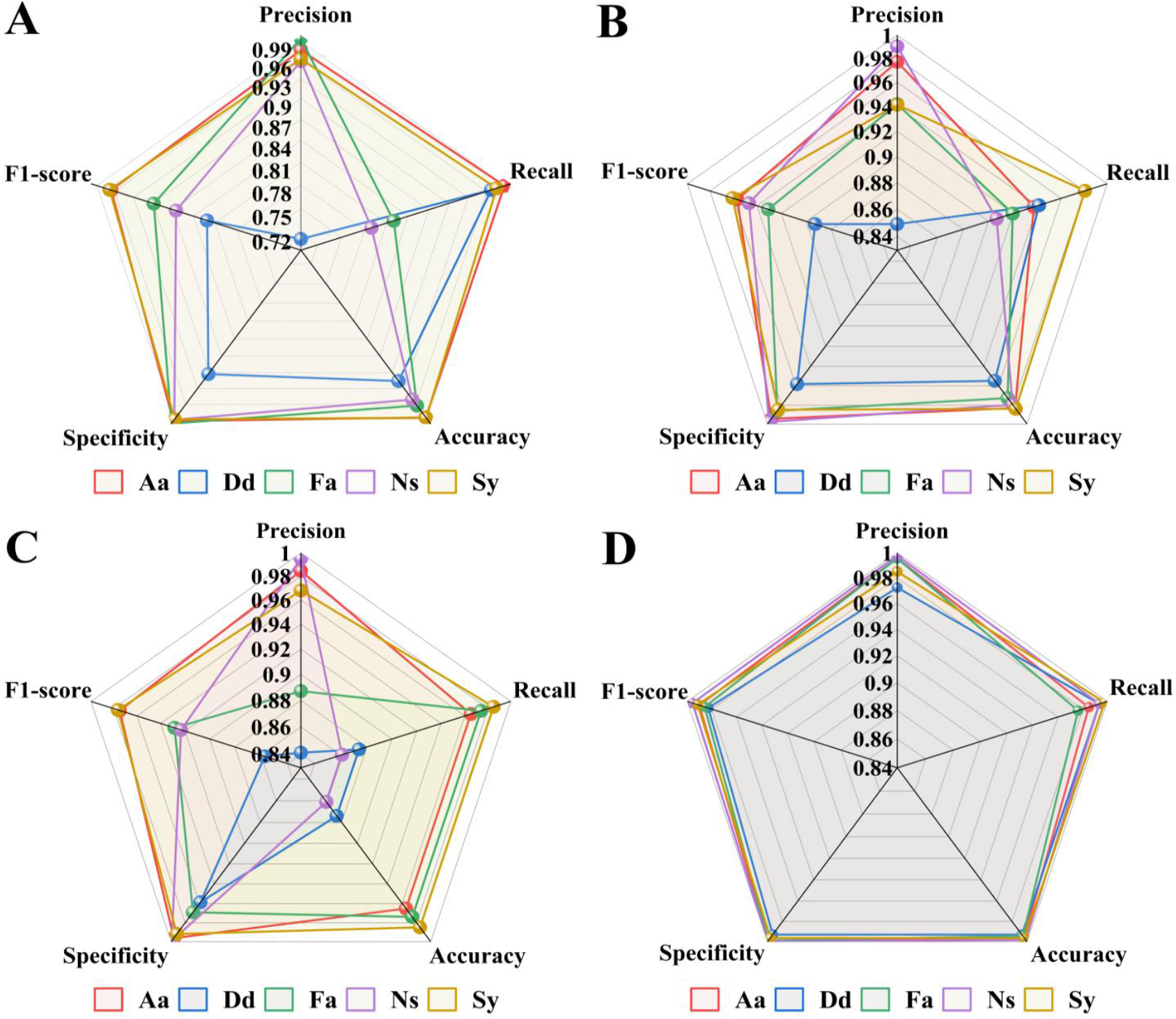
A comparison of assessment metrics across various models. A is the radar chart result of the five evaluation indicators of Improved AlexNet; B is the radar chart result of the five evaluation indicators of Improved AlexNet+MLA; C is the radar chart result of the five evaluation indicators of Improved AlexNet+MSA; D is t he radar chart result of the five evaluation indicators proposed

As shown in Fig. 9, the proposed model outperforms all three compared variants. While the baseline Improved AlexNet exceeds 72% in all categories and the two attention-augmented versions surpass 84%, their performance remains uneven across metrics. In contrast, our model achieves consistently high scores above 97% with minimal inter-metric variance, demonstrating balanced generalization without single-class overfitting. This reliable performance supports its practical utility for unbiased microalgae identification.

A matrix bubble chart is an effective visualization tool that combines a scatter plot layout with variable bubble sizes to represent relationships among three or more dimensions. Fig. 10 employs this chart type to compare the generalization capability of our model against others, plotting their performance in five indicators. In this representation, the size of each bubble is proportional to its corresponding metric value.

**Fig. 10:**
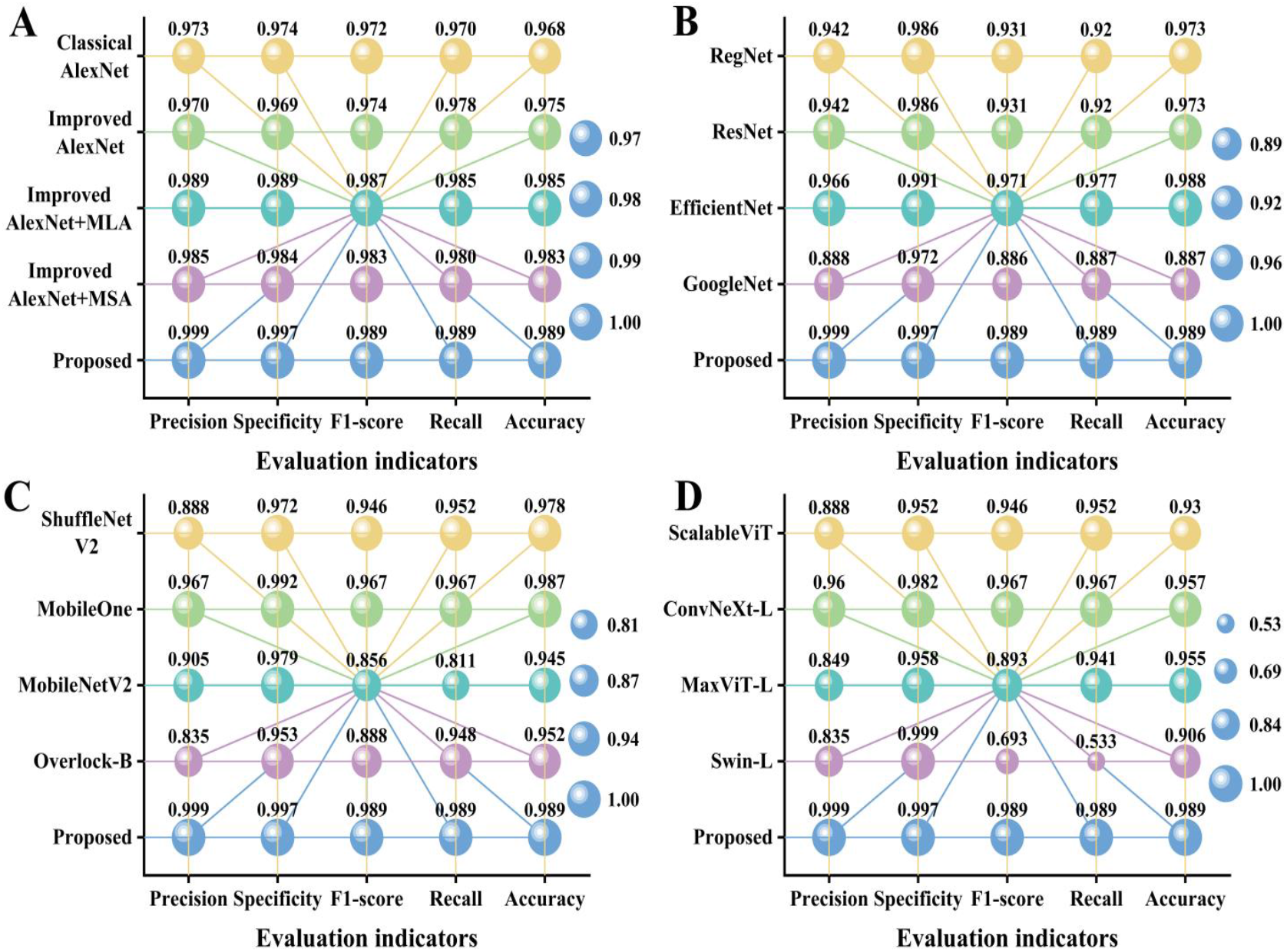
Bubble Chart Comparing Five Indicators of Different Models. A represents different combinations of the methods proposed in this article. B represents the comparison between the model proposed in this article and the basic backbone network. C represents the comparison between the proposed model and lightweight and efficient networks in this article. D represents the comparison between the proposed model and the attention enhanced network.

As illustrated in Fig. 10, the proposed model exhibits a notably larger and more uniformly distributed bubble size compared to other models, indicating stronger and more consistent performance across evaluation metrics. The performance of different architectural combinations varies significantly: both Improved AlexNet+MLA and Improved AlexNet+MSA outperform Classical AlexNet and the baseline Improved AlexNet, confirming the individual efficacy of MLA and MSA in enhancing algae classification. Notably, the full model proposed in this study, which integrates both attention mechanisms, achieves the highest performance among all five combinations. The remaining subFig.s further reveal performance variations across models in identifying different microalgae types. For instance, Swin-L displays uneven bubble sizes, particularly in f1-score and recall, reflecting inconsistent behavior across metrics. In contrast, models such as RegNet, EfficientNet, ShuffleNetV2, and ConvNet-L show more balanced indicator profiles with larger and more uniform bubbles; however, they still fall short of the proposed model, underscoring its superior overall capability.

A grouped 3D scatter plot is used to visualize the relationships among three variables, with data points distinguished by color according to their assigned groups. This representation aids in identifying patterns and clusters across different model categories. For a comprehensive evaluation of classification performance, three representative deep-learning models were selected for comparison. EfficientNet-B0 (Babu et al., 2025) employs a lightweight architecture comprising 16 mobile inverted bottleneck convolutions, along with convolutional, pooling, and classification layers. MobileOne-S0 (Vasu et al., 2023), building on MobileNetV1, integrates RepVGG’s reparameterization concept within MobileOne blocks, balancing accuracy and efficiency via depthwise separable convolutions and a tunable hyperparameter k. OverLoCK-B, a mid-scale variant of the OverLoCK series, enhances long-range dependency modeling and preserves local inductive bias through a top-down attention mechanism combined with hybrid dynamic convolution. These three models represent distinct advanced architectures in deep learning and provide meaningful benchmarks. Their performance relative to the proposed model is depicted in the 3D scatter plot shown in Fig. 11, offering a clear comparative perspective across multiple evaluation dimensions.

**Fig. 11:**
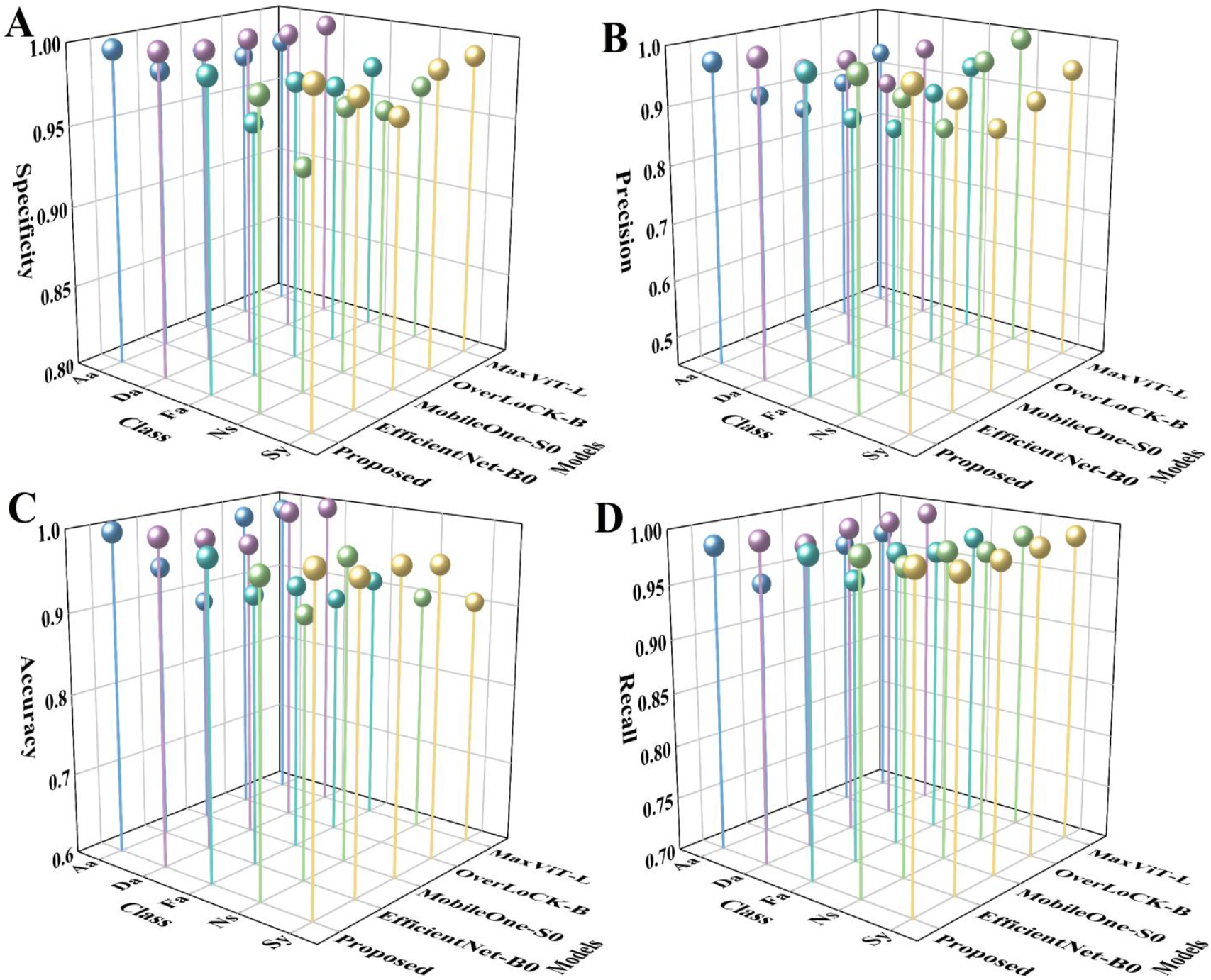
3D scatter plots of different indicators. A is the specificity of four models for each class, B is the precision of four models for each class, C is the accuracy of four models for each class, and D is the recall of four models for each class.

As shown in Fig. 11, the color of the spheres corresponding to each type of microalgae is different, and the height of the spheres is proportional to the value. The higher the sphere, the greater the value. Among the four indicators of Specification, Precision, Accuracy, and Recall, Proposed shows the highest value. Especially in terms of specificity, accuracy, and recall, there are significant differences compared to other models. In terms of accuracy and recall metrics, each model has a high value for the Da category, indicating high performance. But the indicators of the other three models are not balanced among the five types of microalgae classification. Overall, the four higher and more balanced metrics of the model proposed in this article demonstrate its generalization ability in multi class tasks.

In order to compare the probability confidence of the proposed method with other combination methods in the test results and perform a significance test on the mean error of its 95% confidence interval, the results are shown in Fig. 12, where the bar graph represents the mean confidence of each method in identifying each microalgae category, and the error bars on the bar graph represent the 95% confidence interval of the mean.

**Fig. 12:**
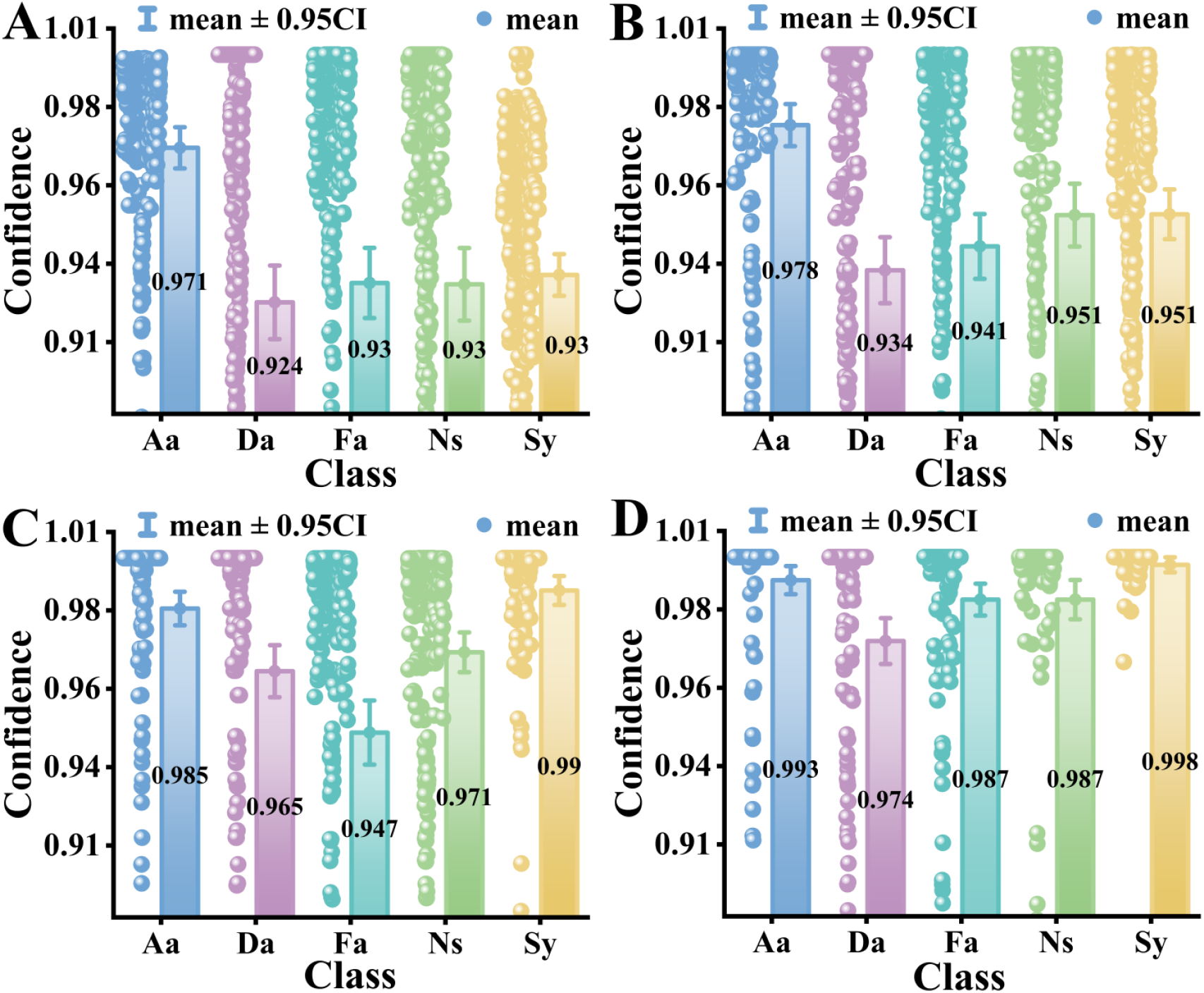
Mean error test results of four methods. A is Improved AlexNet, with a confidence level of over 93% for identifying 5 types of algae; B is Improved AlexNet+MSA, with a recognition confidence level of 93.4% or higher for all five algae species; C is Improved AlexNet+MLA, with a confidence level of over 94.5% for the recognition of 5 algae species; D is Improved AlexNet+MLA+MSA, with a confidence level of 97.4% or higher for identifying 5 types of algae.

Fig. 12 illustrates that Improved AlexNet achieves the highest average confidence for recognizing category Aa, with a confidence interval of [0.98862, 0.99724]; however, its mean confidence across other categories is lower than that of the enhanced variants. In comparison, the incorporation of MSA mechanism significantly elevates the confidence mean for algae recognition across all categories. Conversely, the addition of MLA mechanism markedly narrows the confidence intervals, reflecting improved estimation stability. The combined Improved AlexNet+MLA+MSA architecture attains optimal performance, yielding both a higher average accuracy and a more compact confidence interval than all other methods. Error bar analysis confirms that our proposed combined scheme not only enhances result stability but also meets the criteria for statistical significance in the error bar test, validating its robustness.

A bar chart annotated with significance indicators serves as a fundamental tool for visually assessing module contributions and verifying their effectiveness. It facilitates a direct comparison of the average performance gap between a base model and its enhanced counterparts. By incorporating data point scatter, error bars, and statistical markers, this method distinguishes genuine performance improvements from random fluctuations, thereby providing a reliable measure of both stability and practical significance. Fig. 13 presents a comparison of classification confidence across five model configurations. The vertical axis denotes the confidence level, while the horizontal axis lists the model names. Each bar’s height corresponds to the mean confidence level, with error bars at the top representing the standard deviation (SD), which quantifies data dispersion. Statistical significance between model pairs is indicated using asterisks (* for p ≤ 0.05, ** for p ≤ 0.01, and *** for p ≤ 0.001), offering a clear, quantitative assessment of differential performance.

**Fig. 13:**
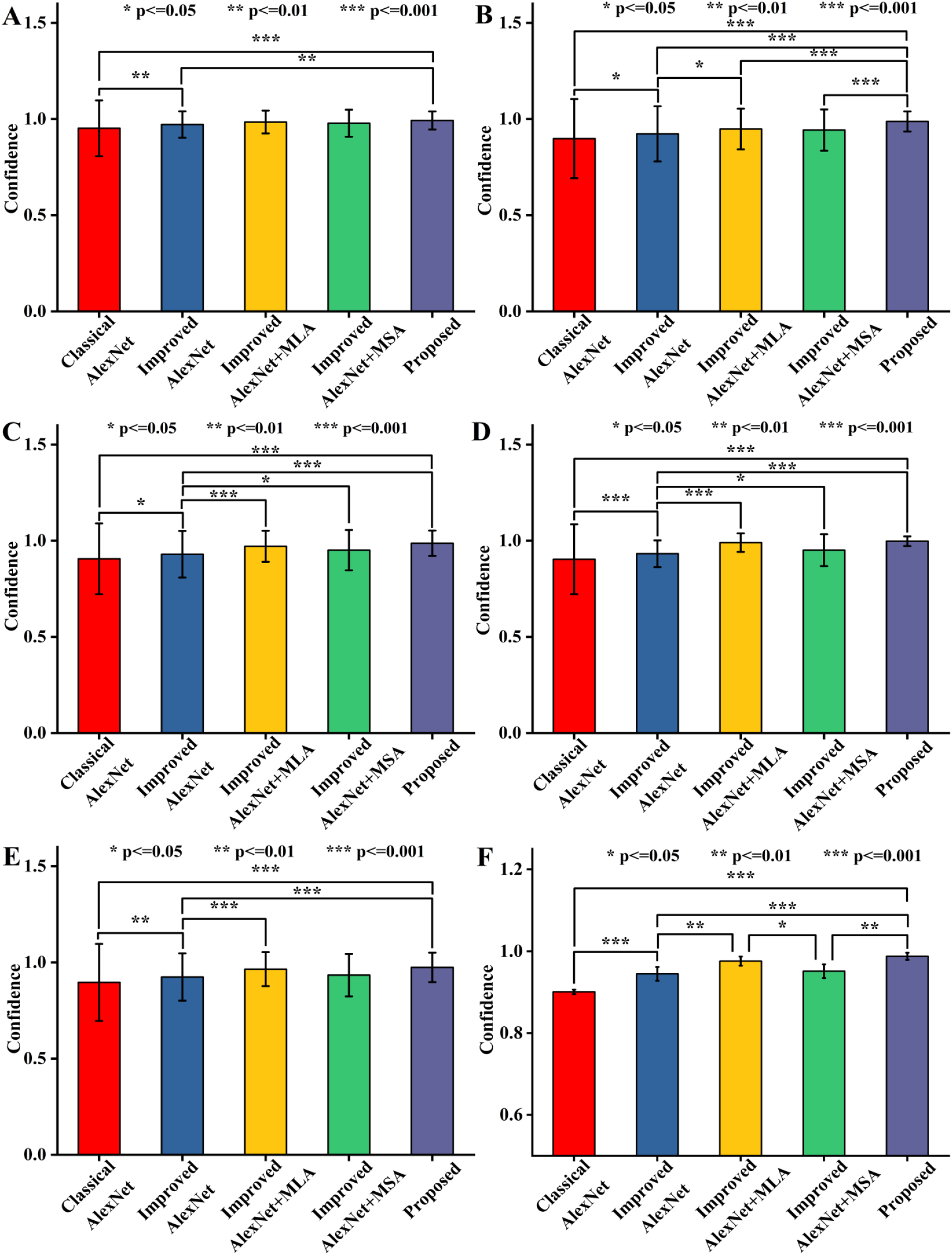
Significant differences in confidence levels among five methods; A is five methods for Aa; B is five methods for Da; C is the five methods for Fa; D is the five methods for Ns; The five methods are Sy’s; F is the confidence level of five methods for all categories.

As raveled in Fig. 13, among the five model configurations compared, the proposed model achieves the highest classification confidence for each sample class, accompanied by the shortest error bars. This result indicates its superior classification capability and reliable output stability. The Improved AlexNet consistently outperforms the Classical AlexNet across all categories, with the difference being particularly significant for class Ns (p ≤ 0.001). This improvement can be attributed to the increased depth of convolutional layers in Improved AlexNet, which enhances its capacity to extract discriminative spectral features. Notably, in subFig. F, all five methods exhibit pairwise statistical significance. Specifically, both Improved AlexNet+MLA and Improved AlexNet+MSA show significant gains over Improved AlexNet alone, confirming that the introduced MLA and MSA modules contribute effectively to performance, thereby validating the feasibility of our architectural strategy. In the preceding subFig.s, the proposed model demonstrates strong statistical significance (p ≤ 0.001) when compared separately to both Classical AlexNet and Improved AlexNet. Overall, the significance testing and error bar analysis consistently affirm that our final model not only achieves substantially better performance than the baseline but also maintains stable and reliable predictions across all evaluated categories.

To visually demonstrate the recognition confidence of the proposed model on the dataset we are using. We randomly select an image from the test set of various algae for testing. The result is shown in Fig. 14.

**Fig. 12:**
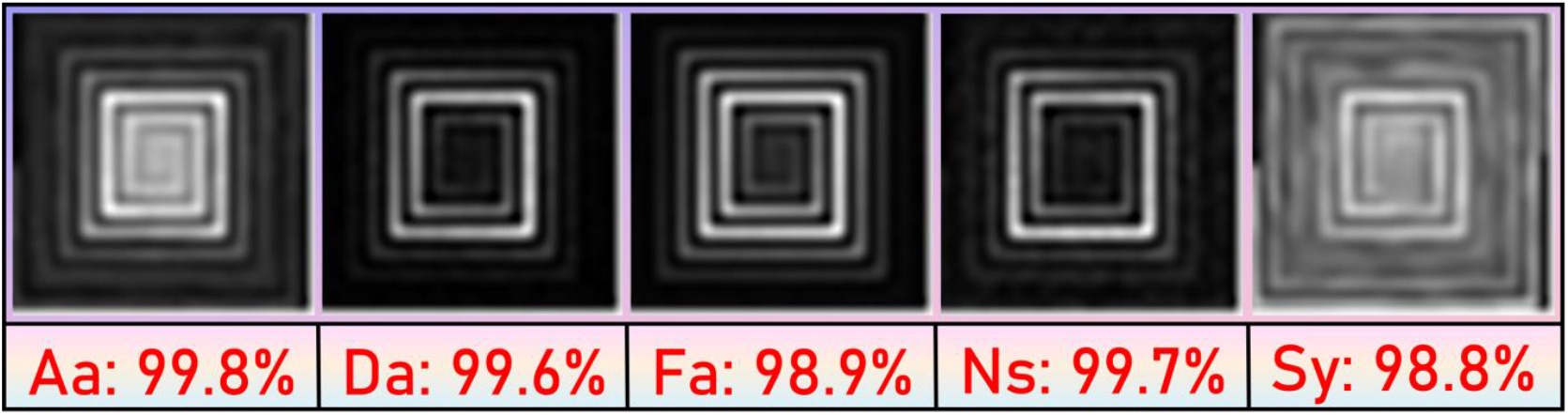
Identification confidence of five types of algae

A precision of 98.91% was obtained on our dataset using an AlexNet architecture with integrated MLA and MSA modules. This model demonstrates superior performance compared to EfficientNet-B0, MobileOne-S0, and OverLoCK-B, alongside a modest parameter count of 16.34M for edge deployment. A benchmarking evaluation was conducted to confirm the inference speed rationale and edge-compatibility of the proposed model. Five optimized models were tested under identical hardware conditions (32 GB RAM, NVIDIA GeForce GTX 1650 Ti), with their FLOPs, operating speed, and accuracy compiled in Table 3.

**Table 3:** Evaluation of Resource Utilization Metrics for Multiple Models.

| Model version. | Params (M) | FLOPs (G) | Speed (s) | Accuracy |
| --- | --- | --- | --- | --- |
| Classical AlexNet | 14.59 | 4.94 | 0.169 | 96.8% |
| Improved AlexNet | 15.36 (↑0.77) | 7.04 (↑2.1) | 0.17 (↑0.001) | 97.45% (↑0.65%) |
| Improved AlexNet+MLA | 16.33 (↑1.74) | 9.63 (↑4.69) | 0.204 (↑0.035) | 98.53% (↑1.73%) |
| Improved AlexNet+MSA | 15.38 (↑0.79) | 7.07 (↑2.13) | 0.183 (↑0.014) | 98.29% (↑1.49%) |
| Improved AlexNet+MLA+MSA | 16.34 (↑1.75) | 9.66 (↑4.72) | 0.211 (↑0.042) | 98.91% (↑2.11%) |

According to the data in Table 3, the improved AlexNet model has increased in both parameter size and floating-point computation compared to the baseline version, but the increase in inference time is limited to only 0.001 seconds. Introducing the MLA module led to a 0.035-second acceleration in inference speed, at the cost of a 1.74M parameter and 4.69G FLOP increase. The integration of the MSA mechanism triggered further growth in parameter count. Despite this increased complexity, experiments confirm that the enhanced AlexNet (with MLA and MSA) strikes a balance on the GTX 1650 Ti, achieving real-time agricultural processing speeds while maintaining moderate resource utilization.This indicates that the model can be effectively deployed on edge devices with limited resources. In the future, through methods such as quantitative compression, inference efficiency can be further optimized to better adapt to low-power embedded platforms.

The approach that is suggested in this study has wide opportunities with regards to edge deployment. The benefits of the AlexNet alone include reduced parameters and reduced computation cost. It is based on this that MSA and MLA modules were added to the AlexNet structure with structural optimization. Despite an increment in the model parameters, the recognition accuracy has been greatly enhanced and the use of resources is at the minimal level. Hence, the model is likely to reach real-time image and video processing and analysis on marginally computing edge devices and low storage capacity. This scheme clearly satisfies the main needs of edge computing in terms of latency and power consumption, and contains a viable technical foundation of the landing of smart edge applications.

## 5. Conclusions

This paper presented ANMM, a lightweight dual-attention neural network for the in-situ hyperspectral classification of marine microalgae. By deepening the AlexNet backbone and strategically integrating MLA and MSA, the model achieves a powerful synergy between local feature refinement and global contextual modeling. An early stopping strategy further ensures robust generalization. Evaluated on a custom dataset of field-collected fluorescence spectra, the model attained a classification accuracy of 98.91%, outperforming several contemporary deep learning models. With a parameter count of 16.34M and low-latency inference, it exhibits strong potential for practical deployment on resource-constrained edge devices within underwater monitoring systems.

The model effectively mitigates overfitting and addresses the challenges of insufficient local detail control and incomplete global feature recognition in processing algal spectral data. However, limitations remain. The dataset’s diversity does not yet cover all relevant marine species and forms. Furthermore, the current system is deployed on personal computers and is not readily transferable to embedded marine monitoring terminals.

To address these issues, future work will collect more microalgae data from diverse marine field environments and employ advanced data augmentation to improve model stability under complex marine conditions. For deployment, the model will be compressed into a lighter version via knowledge distillation for use on low-power marine edge devices.

In summary, the proposed model demonstrates excellent performance and provides a reliable technical framework for autonomous microalgae classification, supporting the development of intelligent ocean observation systems.

## Acknowledgement

This work is supported by National Natural Science Foundation of China (No. 62172140), National Natural Science Foundation of China (No. 81971692), National Natural Science Foundation of China (No. 62176086), Natural Science Foundation of Henan Province (No. 252300421858), Henan Provincial Higher Education Teaching Reform Research and Practice Project under Grant (No. 2024SJGLX0475), Henan Provincial Science and Technology Research Project (No. 242102210032), Henan Province Philosophy and Social Science Education Strong Province Project (No. 2025JYQS0339), and the Scientific Research Fund of Henan University of Urban Construction (No. K-Q2025004).

## Declarations

### Ethical approval/Ethics approval and consent to participate

This article does not contain any studies with human participants or animals performed by any of the authors.

### Informed consent /Consent for publication

This article does not contain any studies with human participants or animals performed by any of the authors.

## Data availability statement

The data and code that support the findings of this study are available from a github website link https://github.com/dyy603/xulaixiang-Microalgae-Spectral-Classification

